# Low-bias RNA sequencing of the HIV-2 genome from blood plamsa

**DOI:** 10.1101/319186

**Authors:** Katherine L. James, Thushan de Silva, Katherine Brown, Hilton Whittle, Stephen Taylor, Gilean McVean, Joakim Esbjörnsson, Sarah L. Rowland-Jones

**Author notes:** These authors contributed equally to the present study (Gilean McVean, Joakim Esbjörnsson, Sarah L. Rowland-Jones). Corresponding authors: Joakim Esbjörnsson, Systems Virology, Lund University, BMC B13, Sölvegatan 19, 221 84 Lund, Sweden,; Sarah L. Rowland-Jones, Nuffield Department of Medicine, NDM Research Building, University of Oxford, Old Road Campus, Headington, OX3 7FZ.

## Abstract

Accurate determination of the genetic diversity present in the HIV quasi-species is critical for the development of a preventative vaccine: in particular, little is known about viral genetic diversity for the second strain of HIV, HIV-2. A better understanding of HIV-2 biology is relevant to the HIV vaccine field because a substantial proportion of infected people experience long-term viral control, and prior HIV-2 infection is associated with slower HIV-1 disease progression in co-infected subjects. The majority of traditional and next generation sequencing methods have relied on target amplification prior to sequencing, introducing biases that may obscure the true signals of diversity in the viral population. Additionally, target-enrichment through PCR requires *a priori* sequence knowledge, which is lacking for HIV-2. Therefore, a target enrichment free method of library preparation would be valuable for the field. We applied an RNA shotgun sequencing (RNA-Seq) method without PCR amplification to cultured viral stocks and patient plasma samples from HIV-2 infected individuals. Libraries generated from total plasma RNA were analysed with a two-step pipeline: (1) *de novo* genome assembly, followed by (2) read re-mapping. By this approach, whole genome sequences were generated with a 28x-67x mean depth of coverage. Assembled reads showed a low level of GC-bias and comparison of the genome diversity on the intra-host level showed low diversity in the accessory gene *vpx* in all patients. Our study demonstrates that RNA-Seq is a feasible full-genome *de novo* sequencing method for blood plasma samples collected from HIV-2 infected individuals.

**IMPORTANCE:** An accurate picture of viral genetic diversity is critical for the development of a globally effective HIV vaccine. However, sequencing strategies are often complicated by target enrichment prior to sequencing, introducing biases that can distort variant frequencies, which are not easily corrected for in downstream analyses. Additionally, detailed *a priori* sequence knowledge is needed to inform robust primer design when employing PCR amplification, a factor that is often lacking when working with tropical diseases localised in developing countries. Previous work has demonstrated that direct RNA shotgun sequencing (RNA-Seq) can be used to circumvent these issues for HCV and Norovirus. We applied shotgun RNA sequencing (RNA-Seq) to total RNA extracted from HIV-2 blood plasma samples, demonstrating the applicability of this technique to HIV-2 and allowing us to generate a dynamic picture of genetic diversity over the whole genome of HIV-2 in the context of low-bias sequencing.

## INTRODUCTION

Human Immunodeficiency Viruses types 1 and 2 (HIV-1 and HIV-2), the two causative viruses of acquired immunodeficiency syndrome (AIDS), are human pathogens of high importance(1). Following the introduction of HIV-1 and HIV-2 into human populations through zoonotic transmission of simian immunodeficiency viruses (SIVs) infecting several species of apes and non-human primates, HIV-1 and HIV-2 are estimated to have infected more than 75 million people worldwide, resulting in over 40 million deaths(2).

Whilst HIV-1 and HIV-2 share some common features, a major difference between the two viruses is the typical viral load associated with chronic infection. In patients infected with HIV-2, viral load is strongly correlated with disease progression and a large proportion (~37% in the Caió cohort) maintained undetectable viral loads and high CD4 counts in the absence of treatment during follow-up (sometimes for more than two decades)(3). Additionally, lack of HIV-2 control is associated with lower viral loads when compared to HIV-1 in patients matched by disease-stage(4-7). Patients with a viral load of more than 10,000 copies/mL can be defined as HIV-2 progressors with a reduced survival probability that is similar to that seen in HIV-1 infected individuals in the absence of treatment(8). HIV-1 disease progression has also been correlated with viral coreceptor use or molecular properties like glycosylation patterns, charge and length of the envelope gene(9-12). Although cytopathic CXCR4 using virions have been isolated from HIV-2 infected individuals in late-stage disease(13, 14), less is known about correlations between molecular properties and disease stage in HIV-2 infection, particularly outside the envelope gene(15, 16). One of the main barriers to a globally protective HIV vaccine is the ability of HIV to evolve rapidly, introducing mutations that abrogate the binding of neutralising antibodies, rendering vaccine responses ineffective(17). Therefore, a major focus of HIV research has been to understand the factors affecting viral evolution and to identify viral epitopes of high conservation as potential vaccine targets(18).

Due to the relatively small size of the full HIV ssRNA genome (~10,000 base pairs [bp]), target enrichment is normally required prior to sequencing in order to generate sufficient DNA for downstream sequencing applications(19). The most common method of target enrichment is PCR amplification(20). This method has two major drawbacks: First, the requirement for detailed *a priori* sequence knowledge to inform robust primer design that ensures the majority of variants in the viral quasi-species are captured(21). Different amplification strategies have shown sensitivities down to 3,000 copies/mL, demonstrating the difficulty of generating robust and high-depth sequence data from patients without detectable plasma viraemia(22, 23). However, the sequence database of HIV-2 is significantly smaller than for HIV-1 and a robust and sensitive pan-HIV-2 primer set has yet to be defined and thoroughly evaluated. Mutations in primer binding sites can also reduce binding efficiency and therefore alter the proportion of specific variants in the final pool of amplicons, or in extreme cases, abrogate primer binding completely, resulting in the loss of that variant in the final analysis(24). Second, PCR is stochastically biased by amplicons from previous cycles acting as templates in the subsequent amplification cycles with the potential to further distort the picture of the viral diversity(25).

Several methods have been proposed to circumvent these problems and reduce the biases introduced into sequencing data through target enrichment. For example, primer-ID allows identification of reads derived from the same viral template through incorporation of a unique eight-mer tag during the reverse transcription of viral RNA(26). Downstream reads can be pooled according to template, and multiple reads from the same template can be used for error correction. A study using Primer-ID observed biased diversity estimates between 2-100-fold when comparing to a library generated without any PCR bias correction, highlighting the importance of considering this factor when sequencing a highly diverse population, such as HIV(26). However, primer-ID still relies on sufficient *a priori* sequence knowledge to allow robust primer design, and the incorporation of the barcode into the 3’ end of the cDNA molecule means it is not applicable to library preparation techniques involving random fragmentation of the target, such as those employed when using Illumina platforms.

Shotgun RNA sequencing (RNA-Seq) has been demonstrated as a powerful tool for the study of RNA viruses(27). Library preparation is performed using random hexamer priming of the total RNA in a sample, negating the need for sequence-specific target enrichment(28). This is particularly desirable for HIV-2, where the sequence data available are significantly limited compared with HIV-1. Few studies have applied RNA-Seq to human RNA viruses. For example, Ninomiya *et al.* applied RNA-Seq to plasma samples taken from two chronically HCV infected patients and demonstrated nearly full-length genome sequences with a mean depth of coverage between 50-70x for the two patients(29). In another study, Batty *et al*. further expanded this method, presenting a high-throughput method for Norovirus sequencing, allowing 77 faecal samples to be sequenced with a mean depth of coverage of 100x and a success rate of more than 99%(30). The authors compared this with a PCR amplification strategy and found that the success rate for whole genome amplification using PCR was 29%. This represents a significant decrease in the performance when compared to RNA-Seq. RNA-Seq has also been used for the discovery of two novel SIVs, demonstrating the power of this method of sequencing without prior sequence information in viral discovery(31).

In the present study, we applied RNA-Seq library preparation methods to both patient plasma samples taken from a rural West African community cohort and cultured lab adapted HIV-2 reference strains. We show that RNA-Seq followed by *de novo* assembly is a feasible and powerful approach when applied to HIV-2 samples with viral loads of at least 5,280 copies/mL. In addition, we demonstrate that RNA-Seq represents a novel, low-bias method of HIV-2 sequencing. Finally, we computed estimates of nucleotide diversity for each gene of HIV-2 on both the intra- and inter-host level. These analyses indicated consistently low estimates of diversity in the accessory gene *vpx* within hosts, highlighting the importance of this HIV-2 specific gene in successful HIV-2 infection.

## MATERIALS AND METHODS

### Patient sample collection and ethics statement

All patient samples used in the present study were collected from members of the Caió community cohort who had provided written and informed consent. Samples were collected prior to the start of the present study. Plasma was separated from whole blood through centrifugation (5000xg, 5 minutes, 4°C) and filtration (0.45μM filter, Millipore, Billerica, MA, USA). Plasma samples were stored at -80°C and transported to Oxford, United Kingdom, in a liquid nitrogen dry shipper. Ethical approval was granted by the Gambian Government/MRC joint ethics committee (#SCC1204) and the Oxford tropical research ethics committee (#170-12).

### In vitro culture of lab adapted HIV-2 reference strains

The lab-adapted HIV-2 strains HIV-2 ROD and HIV-2 CBL20 were propagated *in vitro* in the lymphocyte cell line H9, a single cell clone derived from a HUT 78 cell line. Infection of 5×10^6^ cells was carried out with 200μl of 9×103 TCID/50mL of viral stock. Cells were removed through centrifugation at 250xg for 10 minutes and supernatant was collected on days 3, 5, 7, 9, 11, 13 and 15. HIV-2 concentrations were assayed using the colorimetric Reverse Transcriptase Assay (Roche). For each isolate, the supernatant sample with the highest reverse transcriptase concentration was selected for RNA-Seq.

### RNA extraction, RNA quantification and DNase treatment

Total nucleic acid was extracted from 500μl patient plasma or purified supernatant using the QIAamp UltraSens Viral Kit (Qiagen). Extraction was performed according to the manufacturer’s protocol with the substitution of carrier RNA with linear acrylamide (Ambion) as the nucleic acid co-precipitant. Final elution was performed in 12μl H2O. DNA was removed from the samples through treatment with DNase I (Turbo DNase, Ambion) according to the manufacturer’s protocol. RNA concentration was estimated using the QuBit RNA assay (Invitrogen).

### Library preparation and sequencing

Sequencing libraries were prepared from 5μl of the eluted RNA using the NEBNext^®^ Ultra™ RNA Library Prep Kit for Illumina^®^ (New England Biolabs) according to the manufacturer’s protocol. Sequencing libraries were multiplexed and sequenced using the Illumina HiSeq or MiSeq platforms (Illumina). Patient samples were multiplexed 6/lane (HiSeq), generating 2×100 base pairs paired-end reads and lab adapted strains were multiplexed 2/lane (MiSeq), generating 2×150 bp paired-end reads.

### *De novo* genome assembly and read re-mapping

Sequence data were analysed using a custom pipeline. Reads were trimmed using Sickle, stipulating a median Q-score >30 and a read length >40 bp(32). *De novo* genome assembly was performed using VICUNA(33), with the addition of the optional contamination removal step. During contamination removal, HIV-2 derived reads were identified through similarity to a multiple sequence alignment containing a set of 18 publically available HIV-2 group A sequence data (supplementary information [SI]). Overlapping contiguous sequences generated by VICUNA were assembled into whole genome sequences using the map-to-reference feature in Geneious v6.1.6(34) and manually inspected to derive a whole genome consensus sequence. Consensus genome sequences were manually inspected to ensure that they contained intact open reading frames. Reads were re-mapped to the consensus genome sequence using Bowtie2(35), BWA-SW(36), GSNAP(37) and NovoAlign(38) for each sample. Files containing assembled reads were manipulated using the SAMtools package(39) and downstream statistical analyses and data visualisations were performed using R(40) and the Interactive Genome Viewer(41). Error rates were estimated using the ErrorRatePerCycle feature of GATK(42).

### Quantification of biases

Random hexamer bias was assessed through visualisation of the base composition of reads using FASTQC(43). GC bias was quantified using a custom Python script that scanned the genome using a 50 bp sliding window with a step size of 20 bp. Mean GC content and mean depth of coverage were computed for each window and GC bias was assessed by fitting a linear regression in R(49).

### Analysis of molecular properties

Analyses of molecular properties were performed using an in-house Perl script with potential N-linked glycosylation sites (PNGS) as defined in N-GLYCOSITE(44). Net charge of sequences was determined based on each lysine and arginine contributing +1 and each aspartic acid and glutamic acid contributing -1. Total counts of amino acids were also assessed as described(45). Coreceptor tropism was predicted using four major determinants of dual/CXCR4 coreceptor use (L18Z, V19K/R, V3 net charge >+6, insertions at position 24) (46). CXCR4 use was considered when at least one of the criteria was fulfilled. Sample donors were classified as either having been sampled during the asymptomatic or at AIDS stage (as defined by clinical assessment at the sample time point).

### Phylogenetic analysis

A reference set of 20 HIV-2 group A whole genome sequences were obtained from the Los Alamos HIV Database (Table S1)(44). Reference sequences were aligned with consensus whole genome sequences using Muscle(47) and the alignment was manually inspected using Geneious v6.1.6. A Bayesian phylogeny was inferred using BEAST v1.8.0(48), under the general time reversible model of nucleotide substitution with a proportion of invariant sites and gamma-distributed rate heterogeneity, as determined by jModelTest2(49). The Markov Chain Monte Carlo algorithm was run using 100,000,000 iterations with samples taken from the posterior distribution every 10,000 generations. Following a burn-in corresponding to 10% of the samples, the resulting maximum clade credibility (MCC) tree was visualised using FigTree v 1.4.1(50).

### Estimation of genetic diversity

Mpileup files were generated from assembled reads using the SAMtools package and variants were called using VarScan(51) with a cut-off frequency of 0.05. Nucleotide pairwise diversity (π) was estimated using the Nei and Li method(52) through a custom Python script, taking depth at each position as a proxy for population size and the product of frequency of alternative variants and depth as the number of pairwise differences between sequences. Estimates of diversity were generated for each individual gene and over the whole genome and estimates were normalised using the whole genome average to allow comparison between patients. For comparison, we also calculated diversity on the population level by averaging pairwise phylogenetic tree distances in Garli v2.0(53). This was done for each gene separately based on 200 maximum likelihood bootstrap replicates as described(45).

### Statistics

Two-tailed Fisher’s Exact Test was used to assess categorical data (IBM Corp. IBM SPSS Statistics for Windows, Version 23.0. Armonk, NY: IBM Corp.).

### Nucleotide sequence accession numbers

Nucleotide sequences were deposited in GenBank under the following accession numbers: Will be added before publication.

## RESULTS

### Patient and sample characteristics

Samples from a panel of six members of the Caió HIV-2 community cohort (TD003, TD006, TD013, TD024, TD031, TD062), whose plasma viral loads represented the broad spectrum seen in natural HIV-2 infection, as well as cultures of two lab-adapted HIV-2 strains (HIV-2 ROD and HIV-2 CBL20) were subjected to standard RNA-Seq library preparation (Table 1).

**Table 1.** Clinical data related to the analysed samples and controls^1^.

| Sample | Sampling<br>year | Sex | Country | CD4<br>(cells/μl) | Viral load<br>(cp/ml) | Clinical status<br>at<br>sample date |
| --- | --- | --- | --- | --- | --- | --- |
| TD003 | 2010 | F | Guinea-Bissau | 560 | 82005 | Asymptomatic |
| TD006 | 2010 | F | Guinea-Bissau | 1176 | <50 | Asymptomatic |
| TD013 | 2010 | M | Guinea-Bissau | 509 | 1632 | Asymptomatic |
| TD024 | 2010 | F | Guinea-Bissau | 191 | 10560 | AIDS |
| TD031 | 2010 | F | Guinea-Bissau | 407 | 107183 | Asymptomatic |
| TD062 | 2010 | M | Guinea-Bissau | 497 | 139519 | Asymptomatic |
| CBL20 | 1988 | M | The Gambia | 18 | NA | AIDS |
| ROD | 1985 | M | Cape Verde | 100 | NA | AIDS |

<sup>1</sup>Patients samples: TD003-TD062. Controls: CBL20 and ROD

### Assessment of RNA-Seq using HIV-2 ROD

First, we assessed the performance of RNA-Seq using the well-characterised reference strain HIV-2 ROD. Following initial quality control and removal of low quality reads and adaptor contamination, reads were assessed for the presence of biased random hexamer priming. The remaining high quality reads were assembled to the HIV-2 ROD reference genome sequence (accession number BD413542). The mean depth of coverage over the whole genome was around 2000x for all alignment tools, with GSNAP having the highest mean depth (Table 2). All four alignment tools produced a slightly positive GC bias, and more GC-rich regions tended to have higher coverage. The slopes were very similar (0.79-0.91), implying that the assembly algorithm used does not affect the GC-bias. In order to assess how divergent the HIV-2 ROD that was propagated for the present study was from the published reference sequence, polymorphisms that were fixed at a frequency of >95% in the sample population were annotated as single nucleotide polymorphisms (SNPs) using VarScan (Figure 1). The BWA-SW build was used for this analysis as it agreed with the majority consensus at each site of conflict. All genes except *vif* had SNPs (in total 70 SNPs), and the majority of SNPs were seen in *gag*, *pol* and *nef*. However, when corrected for gene length, *nef* showed the greatest contribution to divergence from the reference genome.

**Figure 1.**
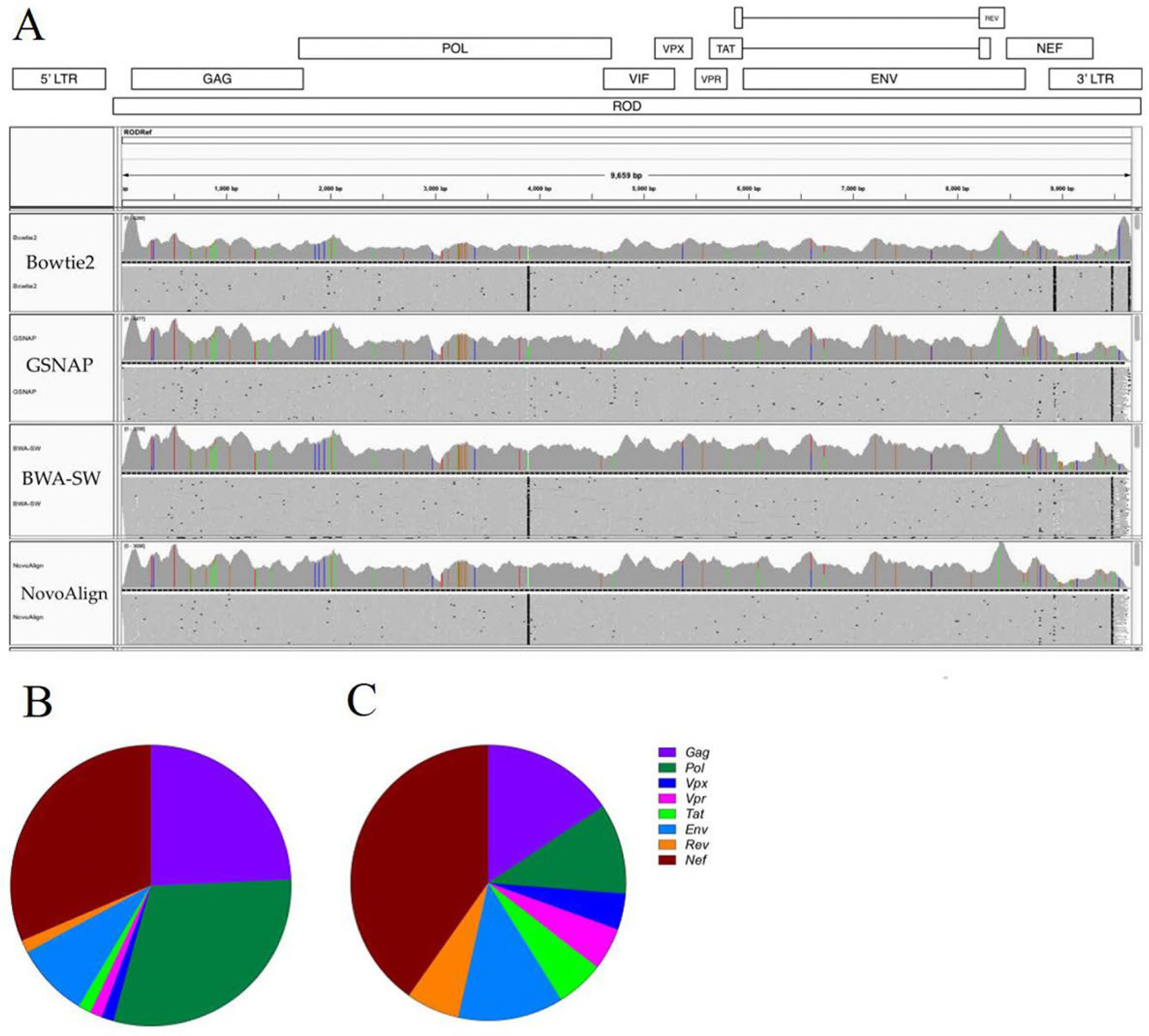
Divergence *in vitro* of the lab adapted HIV-2 isolate HIV-2 ROD. Assembled reads were visualised in Integrative Genomics Viewer (41) and mismatched sites were coloured (A). Sites of conservation with the published reference sequence are shown in grey. Single nucleotide polymorphisms (SNPs) were defined as fixed at a frequency of >95% and the total number of SNPs in each gene were calculated (B). In order to allow for varying gene length, the frequency of SNPs in each gene was also calculated (C).

### *De novo* genome assembly and factors influencing RNA-Seq success rate

After showing that our RNA-Seq approach could be used for whole-genome sequencing of a high-copy number and lab-adapted HIV-2 strain, we assessed the feasibility of using RNA-Seq to generate whole genome sequences directly from primary patient blood plasma samples (Table 1). Clinical blood plasma samples often contain significant amounts of human RNA, making it challenging to perform *de novo* assembly of minority species (such as HIV-2). VICUNA is designed to target populations with high mutation rates and map minority variants into a single consensus sequence, and is therefore particularly suitable for HIV-2, considering the few publicly available HIV-2 whole-genome sequences. In addition, since HIV-2 blood plasma samples usually have significantly lower viral copy numbers than propagated virus isolates, we included a lab-adapted HIV-2 strain derived from a Gambian subject (CBL20), for which the whole-genome sequence is unknown, as a high-viraemic control (Table 1).

In total, whole genome assembly was successful for three of the six patient samples (TD024, TD031, and TD062) and the control (CBL20). Successful patient samples showed complete capture of the coding region of HIV-2 and merged contigs ranged from 9397-9776 bp in length. A merged contig spanning the complete coding region was also assembled for CBL20, demonstrating the applicability of RNA-Seq to both *in vitro* and *ex vivo* samples. These results suggest a cut-off in sensitivity of down to 5,280 copies/mL, with an expectation of at least 0.001% HIV-derived RNA (Table 3). When these limits are considered, the success rate was 75%.

**Table 2.** Summary of read mapping to sample-specific reference sequences.

| **Sample ID** | **Bowtie2** |  | **BWA-SW** |  | **GSNAP** |  | **NovoAlign** |  |
| --- | --- | --- | --- | --- | --- | --- | --- | --- |
|  | **Mean depth** | **Reads aligning** | **Mean depth** | **Reads aligning** | **Mean depth** | **Reads aligning** | **Mean depth** | **Reads aligning** |
| TD024 | 28.53x | 3709 | 31.90x | 3988 | 27.69x | 3426 | 27.79x | 3463 |
| TD031 | 62.33x | 7658 | 67.23x | 8044 | 60.33x | 7172 | 60.50x | 7267 |
| TD062 | 50.01x | 6617 | 59.61x | 7468 | 45.92x | 5658 | 46.64x | 5751 |
| CBL20 | 5502x | 539906 | 4734x | 412557 | 6451x | 618464 | 5156x | 432538 |
| ROD | 1924x | 165506 | 1794x | 152105 | 2146x | 176885 | 1862x | 155696 |

**Table 3.** Samples included in the present study and *de novo* assembly statistics.

| **Sample ID** | **Viral copies<sup>a</sup>** | **Total RNA (ng)<sup>b</sup>** | **Predicted HIV RNA (%)<sup>c</sup>** | **Reads aligning to viral reference** | **Genome covered by all contigs (%)** | **Genes Intact** | **Merged contig length (bp)** |
| --- | --- | --- | --- | --- | --- | --- | --- |
| TD003 | 41002 | 8.70 | 0.0023 | 0 | 0 | 0 | 0 |
| TD006 | <50 | 7.65 | 0.0000033 | 0 | 0 | 0 | 0 |
| TD013 | 816 | 34.00 | 0.000012 | 0 | 0 | 0 | 0 |
| TD024 | 5280 | 2.30 | 0.0011 | 4 998 | 93 | 9 | 9531 |
| TD031 | 53591 | 3.10 | 0.0087 | 9 065 | 90 | 9 | 9397 |
| TD062 | 69759 | 2.85 | 0.012 | 13 304 | 87 | 9 | 9776 |
| CBL20 | >10000000 | 9.65 | >20 | 930 072 | 87 | 9 | 9885 |

<sup>a</sup>Absolute viral input estimated from viral load.
<sup>b</sup>Total RNA input used for library preparation.
<sup>c</sup>Estimated using a viral genome length of 10 kb and absolute viral input.

In order to assess how well *de novo* assembly using VICUNA had captured the HIV-2 genome, consensus sequences were aligned to the commonly used HIV-2 group A reference sequence UC2 (accession number U38293) and annotated according to homology (Figure S1, Table S2). In patient sample TD031, 177 bp of the *gag* leader sequence was missing, whereas the LTR region lacked coverage for patient samples TD024 and TD062, and the reference strain CBL20.

### Phylogenetic analysis of *de novo* genome sequences

A BLAST analysis of the newly generated sequences indicated highest similarity with HIV-2 group A sequences. Bayesian phylogenetic analysis with the 20 publically available HIV-2 group A sequences confirmed this (Figure 2, Table S1). This analysis also showed that the newly generated sequences were clearly distinguishable from existing reference sequences. As expected, the HIV-2 ROD sequence generated in the present study and the published reference sequence were closely related and clustered together with a posterior probability of 1.

**Figure 2.**
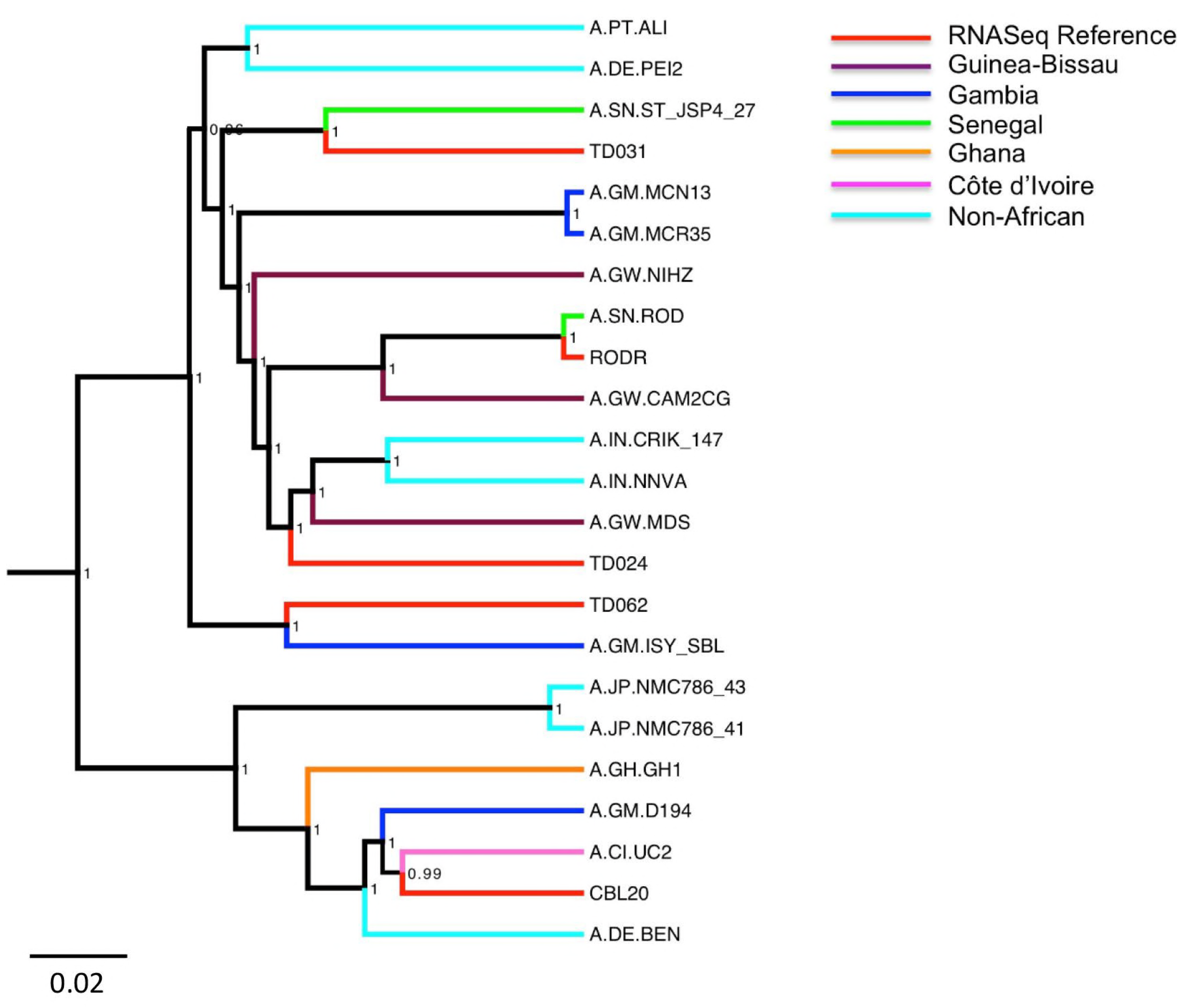
Bayesian phylogeny of HIV-2 genome sequences generated in the present study. Eight-teen whole genome HIV-2 group A sequences were included as a reference set (table S1). Reference sequences are coloured according to country of origin and sequences generated in the present study are shown in red. Bayesian posterior probabilities are included on the corresponding nodes and the scale bar represents the number of nucleotide substitutions per site.

### Read re-mapping to the patient-specific consensus whole genome sequences

In contrast to re-sequencing projects, where a high depth of coverage is required for error correction, deep sequencing of pathogen populations uses high depth of coverage to gain a picture of the diversity in the population as a whole(27). Following *de novo* assembly of a patient-specific consensus genome sequence, we assessed the performance of four commonly used alignment tools when re-mapping reads to the patient-specific consensus (Table 2). Read re-mapping was performed using the total reads without prior HIV-2 enrichment or digital subtraction of human sequences to allow an assessment of how these tools perform in the context of a high level of contamination. This is likely to be a factor of all pathogen sequencing strategies employing RNA-Seq. Mean depth and range of coverage were compared for each aligner (Figure 3). These results show consistent performance of the four aligners, with mean depth of coverage ranging from 28x-67x for the three patient samples. This depth is in line with previous RNA-Seq studies, showing that RNA-Seq is a feasible tool for generating HIV-2 whole genome sequences. Additionally, the high similarity indicates that read mapping is robust and repeatable irrespective of which alignment algorithm that is employed following *de novo* assembly of patient-specific consensus sequences.

**Figure 3.**
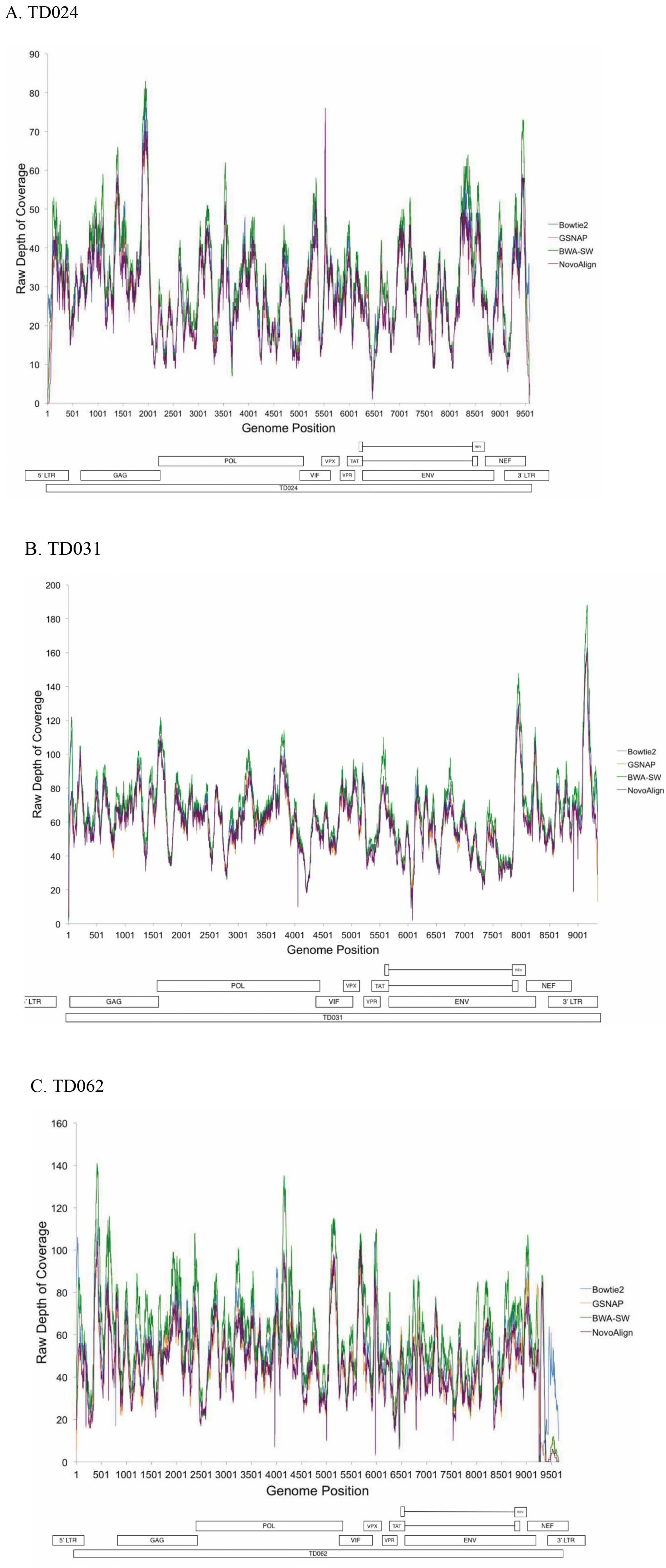
Depth of coverage with the four different aligners. Depth of coverage for each locus was plotted for TD024 (A), TD031 (B) and TD062 (A). Open rectangles represent the locations of HIV-2 genes and the position of the longest merged contig is also shown for each sample. Coverage plots are shown for each of the four aligners, Bowtie2 (blue), GSNP (orange), BWA-SW (green) and NovoAlign (purple). Coverage was plotted as raw depth, showing the number of reads mapping to each locus.

### Assessment of the random hexamer bias

A commonly recognised bias that is specific to RNA-Seq protocols is the random hexamer bias(54). Hypothetical differential binding affinities between different random hexamers result in biased nucleotide composition at the 3’ end of the reads, normally spanning 7-13 bp. Our data indicated that a random hexamer bias affected the first 13 bp of the read (Figure S2). The pattern of the bias was remarkably similar in all three patient samples, suggesting that there may be preferential binding to the same motifs in all samples. This biased read composition can be attributed to random hexamer bias rather than low quality sequencing at the end of the reads as the median Q-score was constant over the length of the read. The effect of the bias did not extend past the first 13 bp of each read and the nucleotide composition stabilised after this point. A correction was not applied to account for the biased nucleotide composition of the first 13 bp, as removal of these positions does not remove the effects of this bias seen in downstream analyses.

### Quantification of the GC% bias and depth of coverage as a function of genomic context

Depth of coverage in samples sequenced using Illumina short read chemistry can be affected by the local GC content of the genome(55). We assessed the effect of local GC content on depth of coverage using a custom script which took a sliding window of 50 bp, with a step size of 20 bp, and calculated GC% and mean depth of coverage in each window. The extent of the GC bias was quantified using the slope of the linear regression line and the bias was assessed for each aligner individually (Table 4). To further compare the different aligners, the mean depth of coverage was normalised in each window using the genome-wide mean depth of coverage (Figure 4). All assemblies showed a slight, positive GC bias, suggesting that GC rich regions had depth of coverage that was higher than the mean. For patient samples TD024 and TD031, the magnitude of the slope was similar for all four aligners, suggesting a constant effect when different assembly algorithms were employed. In contrast, sample TD062 showed more fluctuation between aligners. However, the magnitude of the bias was lowest in this patient, suggesting that the overall effect of the GC bias would be reduced, in spite of the fluctuations. Hence, a positive GC bias in HIV-2 samples sequenced using RNA-Seq may confer variability in depth of coverage over the genome. However, the magnitude of the bias was in line with previous studies and did not show a loss of coverage of any genomic regions due to GC bias.

**Figure 4.**
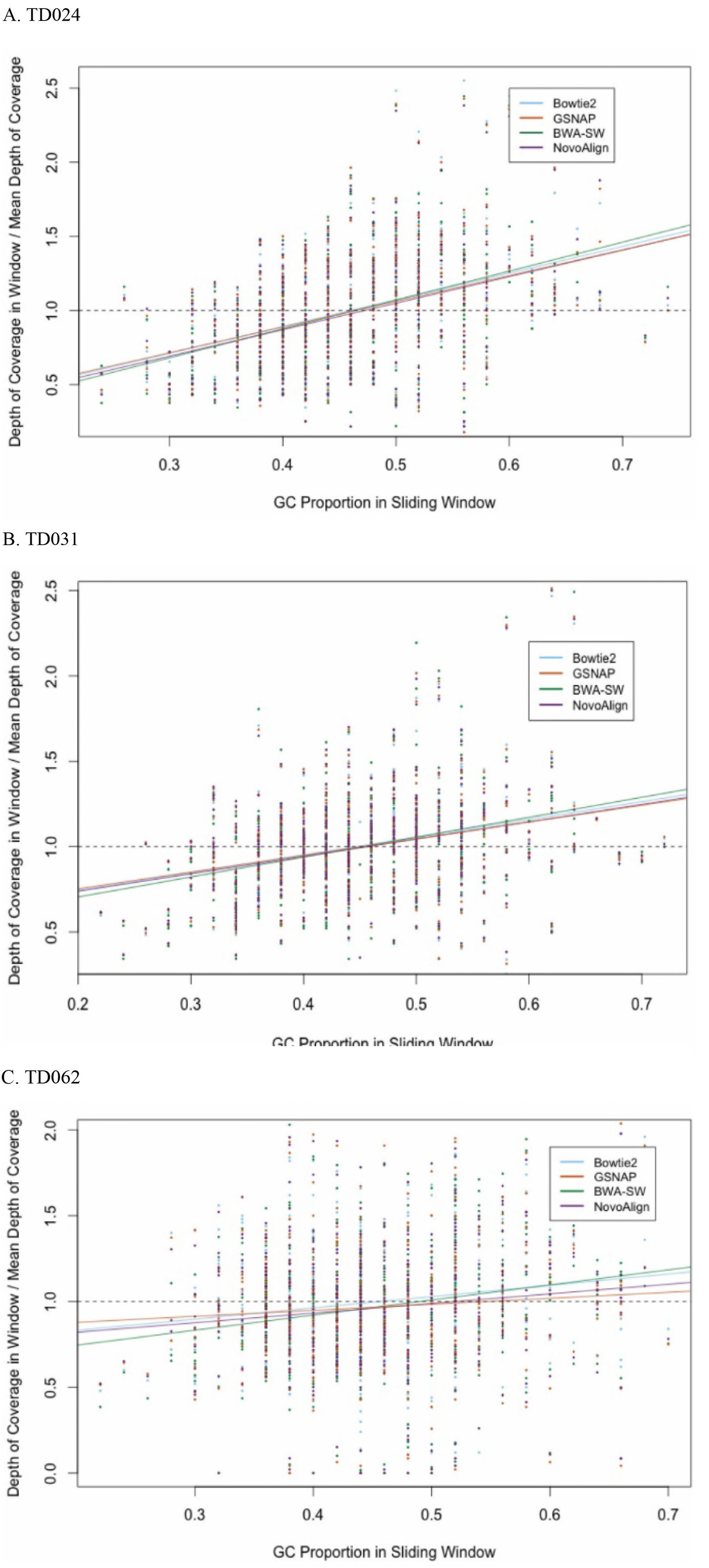
Scatter plots showing the GC bias in assembled reads. GC proportion and normalised depth of coverage in each window were plotted for each aligner individually then grouped by patient sample. Plots are shown for patient TD024 (A), TD031 (B) and TD062 (C). A linear regression was fitted to assess the magnitude and direction of the bias. Regression lines are coloured by aligner. Dashed line indicates the expected regression in the absence of any positive or negative GC bias.

**Table 4.** Summary statistics for the GC% bias present in assembled reads.

| Sample<br>ID | Bowtie2 |  | BWA-SW |  | GSNAP |  | NovoAlign |  |
| --- | --- | --- | --- | --- | --- | --- | --- | --- |
|  | Slope<sup>a</sup> | Intercept<sup>a</sup> | Slope<sup>a</sup> | Intercept<sup>a</sup> | Slope<sup>a</sup> | Intercept<sup>a</sup> | Slope<sup>a</sup> | Intercept<sup>a</sup> |
| TD024 | 1.80 | 0.17 | 1.95 | 0.10 | 1.74 | 0.19 | 1.79 | 0.15 |
| TD031 | 1.04 | 0.54 | 1.17 | 0.47 | 0.97 | 0.56 | 1.02 | 0.54 |
| TD062 | 0.66 | 0.70 | 0.88 | 0.57 | 0.35 | 0.81 | 0.56 | 0.71 |

2 <sup>a</sup>Estimated by fitting a linear regression to the mean values in each sliding window

In order to assess whether genomic context could affect depth of coverage, the HIV-2 genome was partitioned according to gene and mean depth of coverage was compared for each gene individually. The effect of genomic context on depth of coverage was visualised by plotting mean depth of coverage as a function of GC content for each gene (Figure 5). All aligners showed a similar pattern of coverage and no consistent loss of coverage in any genomic region.

**Figure 5.**
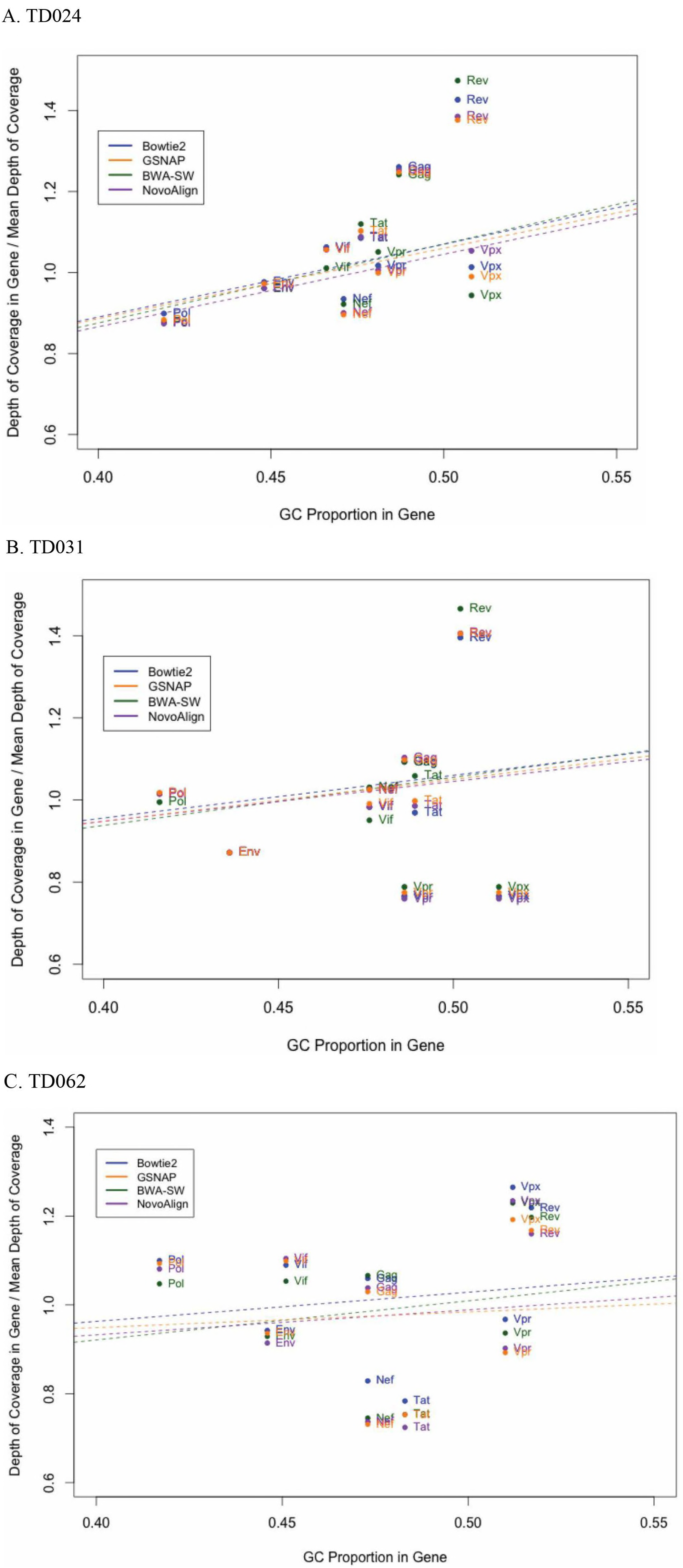
Depth of coverage as a function of genomic context. The depth of coverage was calculated individually for each gene of the HIV-2 genome for TD024 (A), TD031 (B) and TD062 (C). Depth of coverage was coloured by aligner and plotted against the mean GC content of the gene. The predicted GC bias is represented by the dashed line.

### No general trends in molecular properties between the analysed HIV-2 strains or correlations with clinical stage

To characterise the molecular properties of the newly generated sequences and to put them in a broader perspective, we performed an in-depth analysis of these and the 20 selected and publically available HIV-2 group A sequences. Associations between molecular and biological properties were assessed by available clinical and epidemiological data (Table S1). All analyses were performed per HIV-2 gene. In the dataset there were two occasions of duplicate origin, i.e. two sequences that had been generated from the same original patient sample (Table S1: RODR and A.SN.ROD; A.JP.NMC786_41 and A.JP.NMC786_41). These were only counted once when assessing associations between molecular and biological properties between sequences collected during the asymptomatic vs. AIDS stage of disease. No significant differences in sequence length, net charge, total charge, or number of PNGS, in any of the nine HIV-2 genes, were found between sequences from asymptomatic (N=6) and AIDS stage patients (N=12, Table S1). Prediction of coreceptor tropism based on the *env gp120 V3* region indicated that 50% (three of six) and 58% (seven of 12) of the participants had CXCR4 using viruses in the asymptomatic and AIDS stages, respectively (p=1.00, two-tailed Fisher’s Exact Test, Table S3). Furthermore, we found no diagnostic motifs or amino acids between asymptomatic and AIDS stage patients in any of the nine HIV-2 genes (Figures S3).

### Genome-wide estimation of genetic diversity in HIV-2 in the context of low-bias sequencing

To determine how the diversity varies over the HIV-2 genome, we estimated nucleotide pairwise diversity from assembled reads (Bowtie2 assembly) using a custom script. Raw estimates of diversity of the whole genome was 0.0010 substitutions/site for TD024, 0.0007 substitutions/site for TD031, and 0.0014 substitutions/site for TD062. In comparison, the raw estimates for the *env* gene was 0.0013 substitutions/site for TD024, 0.0008 substitutions/site for TD031, and 0.0020 substitutions/site for TD062. To compare the relative genetic diversity between the patients, we normalized the raw estimates using the genome average (Figure 6). Overall, our analysis showed similar results between patients with the highest level of within-host diversity seen in *env* for all three patients, whereas the lowest diversity was seen in the *vpx* and *rev* genes. Interestingly, the diversity in *pol* seemed to be higher than for the genes *gag, vpx*, *tat, rev* and *vif* for all three patients.

**Figure 6.**
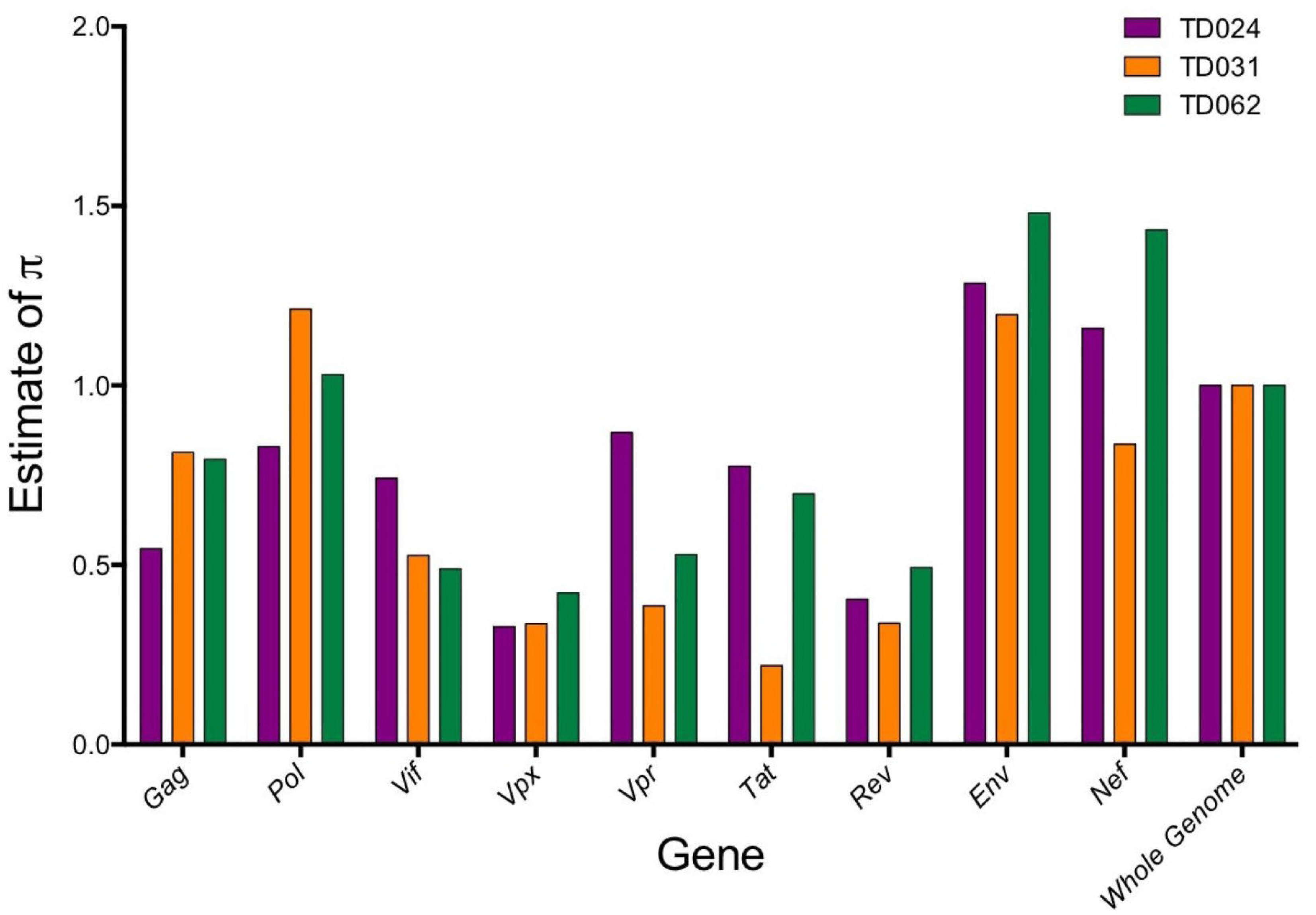
Nucleotide diversity plot. Nucleotide site diversity estimates per gene, normalised to the whole genome estimate. Diversity was estimated for samples TD024 (purple), TD031 (orange) and TD062 (green). Calculating the diversity relative to the whole genome estimate as performed to allow a comparison between patients.

To compare the above results of intra-host viral diversity in different HIV-2 genes with viral diversity in different HIV-2 genes *between* hosts, we performed a phylogenetic bootstrap analysis of our newly generated whole-genome sequences and the reference sequences. This analysis showed that, similarly to the intra-host viral diversity above, the *env* gene was the most diverse gene, followed by the *nef* gene. However, in contrast to the intra-host analysis, this analysis indicated that *pol* was the second least diverse gene when compared between hosts (only *vif* was less diverse, Figure S4).

## DISCUSSION

Deep sequencing of HIV offers unparalleled opportunities to gain a high-resolution picture of the nature and diversity of the viral quasi-species in a single patient. Our study presents a novel and robust pan-HIV-2 whole genome amplification strategy using RNA-Seq, allowing the entire coding region of HIV-2 to be sequenced without the need for detailed *a priori* sequence knowledge. We show a broad applicability of this method, presenting data from both lab-adapted isolates and patient plasma samples. To our knowledge, only one previous study has used a next-generation sequencing approach to determine the full-genome of HIV-2(56). However, we used HIV-2 isolates propagated in cell culture prior to library preparation, and aligned the generated sequence reads to a common reference strain (HIV-2 BEN). We analysed patient samples and demonstrated a cut-off in sensitivity down to a viral load of 5,280 copies/mL, and an expectation of at least 0.001% HIV-2 RNA in the sample. When these conditions were fulfilled, we report a success rate of 75%, which is lower than previously reported by Batty *et al.* when applying RNA-Seq to Norovirus. However, the lower HIV-2 plasma viral loads of the patient samples used in the present study readily explain this reduced success rate. A cut-off of 5,280 copies/mL restricts this method to viraemic HIV-2 patients, and it is possible that an alternative approach would be needed for samples with lower viral loads. However, we anticipate that RNA-Seq could also be successfully applied to samples taken from untreated HIV-1 patients, where the typical viral load is 10 to 1000 times higher than for HIV-2.

Whilst RNA-Seq allows whole genome sequencing of HIV-2 without the need for detailed sequence knowledge, the lack of sequence-specific target amplification also leads to a reduction in the use of PCR amplification and the resulting biases, generating sequence data that is more representative of the true population frequencies. Here, we aimed to quantify the other biases known to be associated with RNA-Seq. We found evidence of a moderate positive GC bias, which varied between samples but was consistent when different aligners were used. We also found evidence of a biased nucleotide composition in the first 13 bp of the reads, suggesting the presence of non-random random hexamer priming. Although these biases could be responsible for the fluctuations in coverage over the genome, we observed no correlation between genomic location and depth of coverage. This suggests that these fluctuations were randomly distributed and not due to the varying diversity seen in different functional genomic sites.

Whilst patient consensus sequences contained all nine genes of HIV-2 in intact reading frames, there was some variability in the assembly of the 3’ and 5’ LTR and the *gag* leader sequence. In patient sample TD031, the loss of 177 bp of the *gag* leader sequence can be attributed to the failure of the RNA-Seq library preparation method to capture this region. The initial fragmentation step in library preparation can lead to the loss of distal regions of the RNA molecule and this is the most probable cause of the lack of coverage in this genomic region. For patient samples TD024 and TD062 and the reference strain CBL20, the lack of coverage was probably due to the nature of the LTRs in HIV-2. The 5’ and 3’ LTR regions only exist as true 990 bp repeats in the proviral form of the virus, whereas in the RNA genome, the 5’ LTR comprises the R and U5 regions and the 3’ LTR is composed of the R and U3 regions(57). The sequence alignment used during assembly contained HIV-2 sequences from both cDNA and RNA HIV-2 genomes, assembly was conducted using ‘complete’ LTRs, both containing U5, R and U3(44). Ambiguous read mapping is normally resolved by using the location of the read mate to provide information on the most likely coordinates. In the present study the insert size (250-350 bp) and the nature of the LTRs meant problematic mapping in the case of reads mapping to the R region, as the read mate will also fall in the LTR. Therefore, it was not possible to resolve the correct orientation of the reads, resulting in the loss of coverage from one LTR.

The ability to sequence the whole HIV-2 genome in a single experiment allowed us to compare pairwise nucleotide site diversity between the different genes of HIV-2. In addition, it has been suggested that HIV-1 genetic diversity was reduced in the context of HIV-1 and HIV-2 dual infection, when compared to matched individuals who had HIV-1 mono-infection(58). Although HIV-2 genetic diversity was not examined in that study, the reduced rate of disease progression for HIV dual-infected individuals suggests that there may be an epistatic interaction in the context of HIV dual-infection. The current study showed similar patterns of within-host diversity across all three patients. The highest diversity was seen in *env*, an observation that is in line with patterns seen in HIV-1. HIV-2 partial *env* diversity has been estimated through different approaches in previous studies, and, although on the lower side, our estimated intrahost *env* diversities were in the same range as previous shown in previous studies that used molecular cloning for sequence generation(15, 59, 60). It is likely that the high level of diversity in *env* is largely driven by selective pressures of the host immune system, driving escape of immune responses. Similarly, a high level of diversity was seen in *nef* in all three patients. *Nef* is an accessory gene that has a key role in the evasion of host immune responses, primarily through HLA and CD4 down-regulation, preventing the display and recognition of virally derived peptides at the cell surface. Presumably the high diversity in HIV-2 *nef* can be tolerated without significant loss of Nef function or reduced viral fitness. In HIV-1 infection, *pol* is thought to be highly conserved for functional reasons and therefore typically shows a relatively lower diversity compared with for example the *env* gene(61). In contrast, we observed a high level of within-host HIV-2 *pol* diversity in all the subjects studied here. Interestingly, a recent study has shown a high level of within-host diversity in *pol* following vertical HIV-1 transmission(62). Although we do not know the route of transmission of our study subjects, it is likely to be horizontal, and it is tempting to speculate that there may similarities between HIV-1 vertical transmission and horizontal HIV-2 transmission. That is, in contrast to horizontal HIV-1 transmission where the majority of infections are attributable to a single transmitted-founder virus(63), multiple transmitted founder viruses may cause the majority of HIV-2 infection. This remains, however, to be determined in future studies on acute HIV-2 infection (which has been difficult to capture as indicated by only one described adult case of acute HIV-2 infection)(64). Some potential caveats of our intra-host diversity analysis exists:

(1) HIV-2 diversity has been reported to increase over the course of infection(15, 59). It is possible that parameters like the duration of infection or the mode of transmission influenced the diversity level. Lack of such information combined with diversity estimates from only three patients prevented us from analyzing such associations. (2) For some single nucleotides over the genome, the coverage was lower than 20 sequence reads, and from a sample-perspective, the depth of coverage was positively correlated with the viral copy number. On the one hand, low coverage may underestimate the true genetic diversity. On the other hand, some regions of the genome are evolutionary conserved, and there is a limited number of virus variants that can theoretically co-exist in a sample with low viral load. However, there are limited data about the nature of genetic diversity in HIV-2. A detailed understanding of virus diversity has implications for vaccine design, development of drug resistance and disease pathogenesis.

*Vpx* is an HIV-2 specific accessory gene that is entirely absent from the HIV-1/SIVcpz lineage. The role of *vpx* is antagonism of the host restriction factor SAMHD1, which blocks reverse transcription of viral RNA in slowly dividing cells such as macrophages and resting CD4+ T cells(65). However, little is known about the implications of this antagonism on the course of HIV-2 disease progression. The observation of a consistently low level of diversity in *vpx* may be indicative of a high level of conservation in *vpx*, suggesting that *vpx* has a critical role in the maintenance of high level of HIV-2 viraemia. The different roles of *vpx* in HIV-2 infection remain to be clearly defined, but a recent study by Yu *et al*. identified a SNP in a *vpx* allele derived from a viraemic patient that totally abrogated the ability of *vpx* to promote SAMHD1 degradation *in vitro*(66).

In conclusion, we show that RNA-Seq library preparation methods can be applied to HIV-2 blood plasma samples. Resulting *de novo* genome assemblies captured the entire coding region of HIV-2 in intact open reading frames and read re-mapping allowed us to demonstrate the importance of a two-step analysis pipeline. In the context of a highly diverse retrovirus, such as HIV-2, the selection or generation of an appropriate reference sequence is a critical first step, allowing robust and repeatable down-stream read mapping. We also demonstrated a low level of GC and random hexamer bias, and in the absence of sequence-specific target amplification, show that RNA-Seq offers a method of whole genome HIV-2 sequencing in a low bias context. However, some challenges in RNA-Seq remain. For example, although the sequencing costs have fallen dramatically in recent years, RNA-Seq is still expensive and costs continue to be a barrier to an even more widespread adoption. In the present study, we multiplexed six patient samples using the Illumina HiSeq in order to reach a mean depth of coverage of up to 67x. Although this coverage is more than sufficient for consensus sequence calling, it may have be too low if the primary goal was to determine minority variants (at least in samples with high viral loads). Hence, the importance of developing novel and low-bias HIV sequencing protocols cannot be understated, as the ability to gain a complete and accurate picture of HIV genetic diversity is critical to the development of globally effective and preventative HIV vaccines.

## Acknowledgements

KJ was supported by a Wellcome Trust 4-year PhD-Studentship (Grant No. H5RSZMO). JE was supported by the Swedish Research Council (350-2012-6628 and 2016-01417) and the Swedish Society of Medical Research (SA-2016). We thank Shokouh Makvandi-Nejad and Lorna Witty for their input into the project design and Takayuki Chikata and Masafumi Takiguchi for their sequencing support.

